# Morphological and ecological notes on the swimming crab *Achelous hastatus* (Linnaeus, 1767) (Family: *Portunidae*), from Tenerife, Canary Islands (NE Atlantic)

**DOI:** 10.1101/2025.08.11.669774

**Authors:** Michael Bommerer

**Affiliations:** Institute for Molluscan Systematics. Research Group Swimming Crabs (Family Portunidae), Tenerife, Canary Islands, Spain

**Keywords:** Achelous hastatus, Portunidae, morphometrics, voucher specimen, Canary Islands, Macaronesia, OBIS, GBIF, nocturnal foraging, scavenging behaviour, marine biodiversity, Atlantic–Mediterranean distribution

## Abstract

A voucher-backed adult male specimen of *Achelous hastatus* (Linnaeus, 1767) was collected on 6 August 2025 during a night dive at 6 m depth off Granadilla, Tenerife, Canary Islands. The specimen was subjected to comprehensive morphometric analysis and documented with high-resolution imagery under controlled laboratory conditions. Morphological features—including a broadly hexagonal carapace with nine anterolateral teeth (ninth markedly elongate), a four-lobed frontal margin, robust chelipeds with slender fingers, and a natatory fifth pereiopod—matched diagnostic descriptions from comparative literature. Field observations documented nocturnal foraging in association with a multi-individual feeding aggregation, scavenging on vertebrate remains. Aggregated occurrence data from the Ocean Biodiversity Information System (OBIS) and the Global Biodiversity Information Facility (GBIF) confirm the species’ widespread Atlantic–Mediterranean distribution, with Spain (including the Canary Islands) representing a primary stronghold. This record contributes high-resolution imagery, morphometric detail, and ecological context to the biogeographic dataset for *A. hastatus* in the Macaronesian region.

## 1. Introduction

*Achelous hastatus* (Linnaeus, 1767) is a marine brachyuran crab in the family *Portunidae* Rafinesque, 1815, subfamily *Achelouinae* Spiridonov, Neretina & Schepetov, 2014 (superfamily *Portunoidea* Rafinesque, 1815). Originally described by Linnaeus as *Cancer hastatus*, the species has undergone multiple reassignments—appearing historically under combinations such as *Neptunus hastatus, Portunus hastatus*, and *Lupea hastata*—before its current placement in the genus *Achelous* De Haan, 1833 (Rodrigues et al., 2017; Spiridonov et al., 2014; DecaNet eds., 2025).

Its geographic distribution spans tropical and subtropical regions of both the eastern and western Atlantic, including the Mediterranean Sea and Caribbean Sea, and along the coasts of West Africa, Cape Verde, the Azores, Madeira, and the Canary Islands (Rodrigues et al., 2017; DecaNet eds., 2025). According to GBIF occurrence data (GBIF.org, 2025), A. hastatus is represented by 958 georeferenced records worldwide. The majority are from Spain (657 records, including 24 from the Canary Islands), followed by Italy (65), France (55), Portugal (16), and Lebanon (14). Remaining records are distributed among other coastal nations within the species’ known Atlantic-Mediterranean range. According to aggregated occurrence data from the Ocean Biodiversity Information System (OBIS, 2025), A. hastatus is represented by 52 georeferenced records compiled from nine contributing datasets, encompassing citizen science observations, institutional collections, and historical expeditions. The majority of records are from Spain (36), followed by Greece (2), Egypt (4), Israel (1), Tunisia (1), and Morocco (1). The remaining six records lack explicit country attribution but include two from Gran Canaria (Canary Islands), two from Crete (Greece), and one from the South Atlantic. The OBIS dataset integrates contemporary contributions such as the Seawatchers Marine Citizen Science Platform (Chic & Garrabou, 2020) and Programa Poseidon (ECOAQUA Institute, 2015), as well as legacy holdings from museum collections (Olivas González, 2016; Santos, 2016) and historical faunal surveys near Alexandria, Egypt (Balss, 1936). Ecologically, *A. hastatus* inhabits shallow marine habitats with mixed sand–rock substrates, seagrass meadows, and coastal reef zones. As a strong swimmer, it utilizes its flattened fifth pereopods as natatory paddles and exhibits scavenging and predatory feeding, active both during the day and night (Rodrigues et al., 2017). Known depth records range from the intertidal zone to ∼60 m, though most observations are from depths under 20 m.

Despite its regional presence, high-resolution photographic documentation and precise morphometrics from the Canary Islands remain limited. This study aims to address that gap by reporting a voucher-backed male specimen collected at 6 m depth off Granadilla, Tenerife, on 6 August 2025. The specimen is documented with standardized dorsal and ventral photography and comprehensive morphometric measurements following portunid protocols. This contribution enhances the taxonomic and morphological dataset for Achelous in the northeastern Atlantic and furnishes a robust baseline for future taxonomic, ecological, and biogeographic research in the Macaronesian region.

## 2. Materials and Methods

### 2.1 Specimen Collection and Habitat

An adult male specimen of *Achelous hastatus* was collected on 6 August 2025 during a survey night dive at 01:00 local time off Granadilla, Tenerife, Canary Islands (28°05′06″ N, 16°29′23″ W). The specimen was encountered at a depth of approximately 6 m within a sandy patch of ∼100 m^2^ adjacent to mixed rocky substrate and the marina walls. It was part of an estimated aggregation of ten or more conspecifics, all actively feeding on organic remains from barbecue pork bones dispersed across the substrate. The dive had a total duration of 113 min, with a maximum depth of 8 m, using a single 11 L, 230 bar compressed-air cylinder. The specimen was captured by hand and subsequently preserved as a voucher.

### 2.2 Photography

The specimen was transferred to a laboratory facility and photographed in dorsal and ventral views under controlled lighting conditions using a high-resolution digital camera (Canon 80D). A millimeter scale bar was included in each image for size reference. One dorsal-view image (Figure B-1) and one ventral-view image (Figure B-2) were captured to document the general morphology. Image contrast and sharpness were adjusted using Adobe software to improve visual clarity without altering morphological details.

**Figure B_1.**
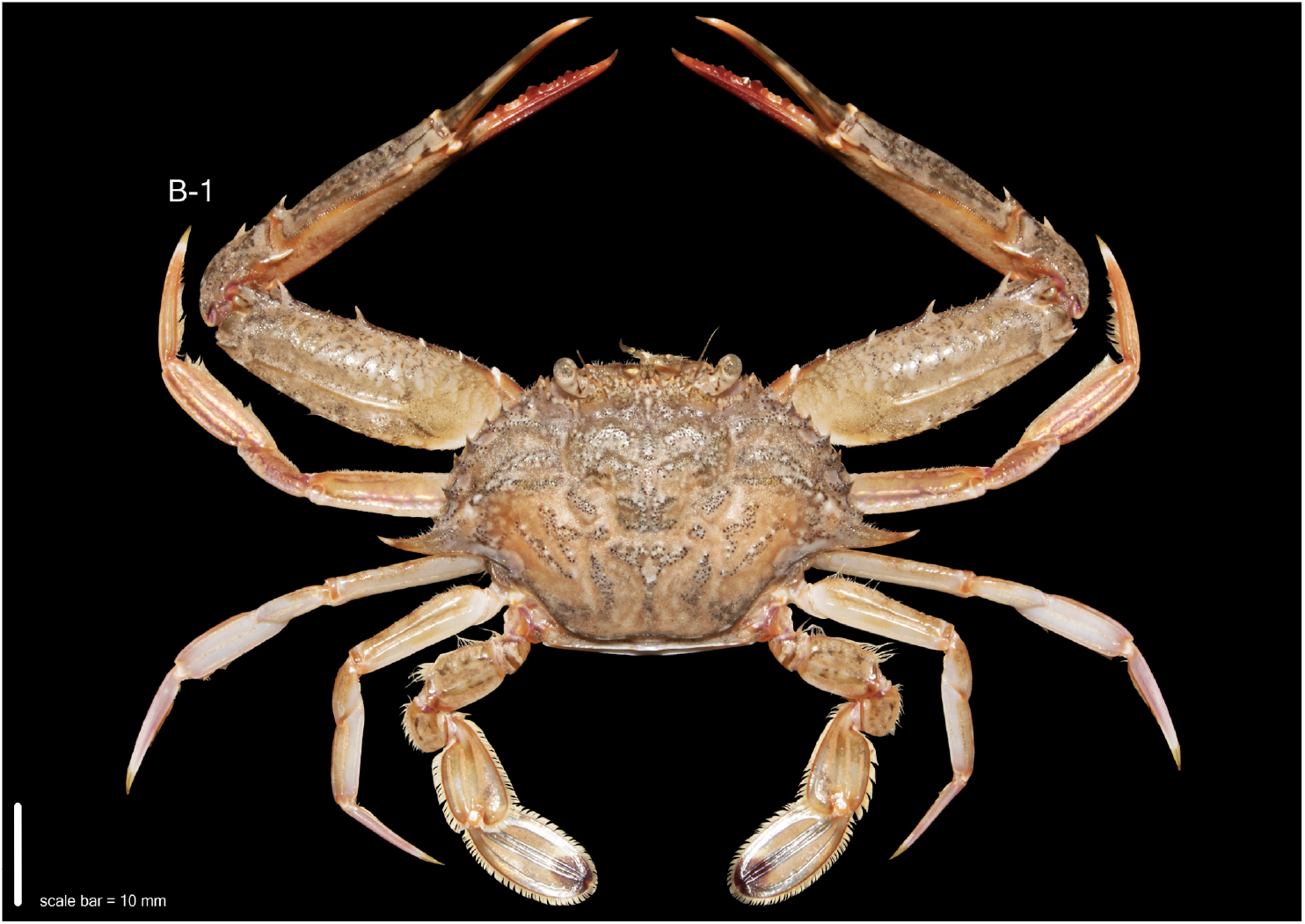
Dorsal view of *Achelous hastatus* (Linnaeus, 1767) (*Decapoda*: *Brachyura*: *Portunidae*: *Achelouinae*), adult male voucher specimen (Specimen ID: IMS-AHA-001) collected from Granadilla, Tenerife, Canary Islands (28°05′06″ N, 16°29′23″ W). The specimen was encountered at a depth of 6 m in a sandy patch adjacent to mixed rocky substrate and marina walls during a night dive on 6 August 2025. Photographed under standardized laboratory conditions. Morphometric measurements: carapace width (CW) = 48.2 mm; carapace length (CL) = 23.1 mm; frontal width (FW) = 8.7 mm; frontal spine width = 1.0 mm; cheliped total length = 74.7 mm; propodus length (cheliped) = 39.5 mm; fixed finger length = 39.5 mm; dactylus length (cheliped) = 16.8 mm. Scale bar = 10 mm. Associated iNaturalist record: https://www.inaturalist.org/observations/304259129

**Figure B_2.**
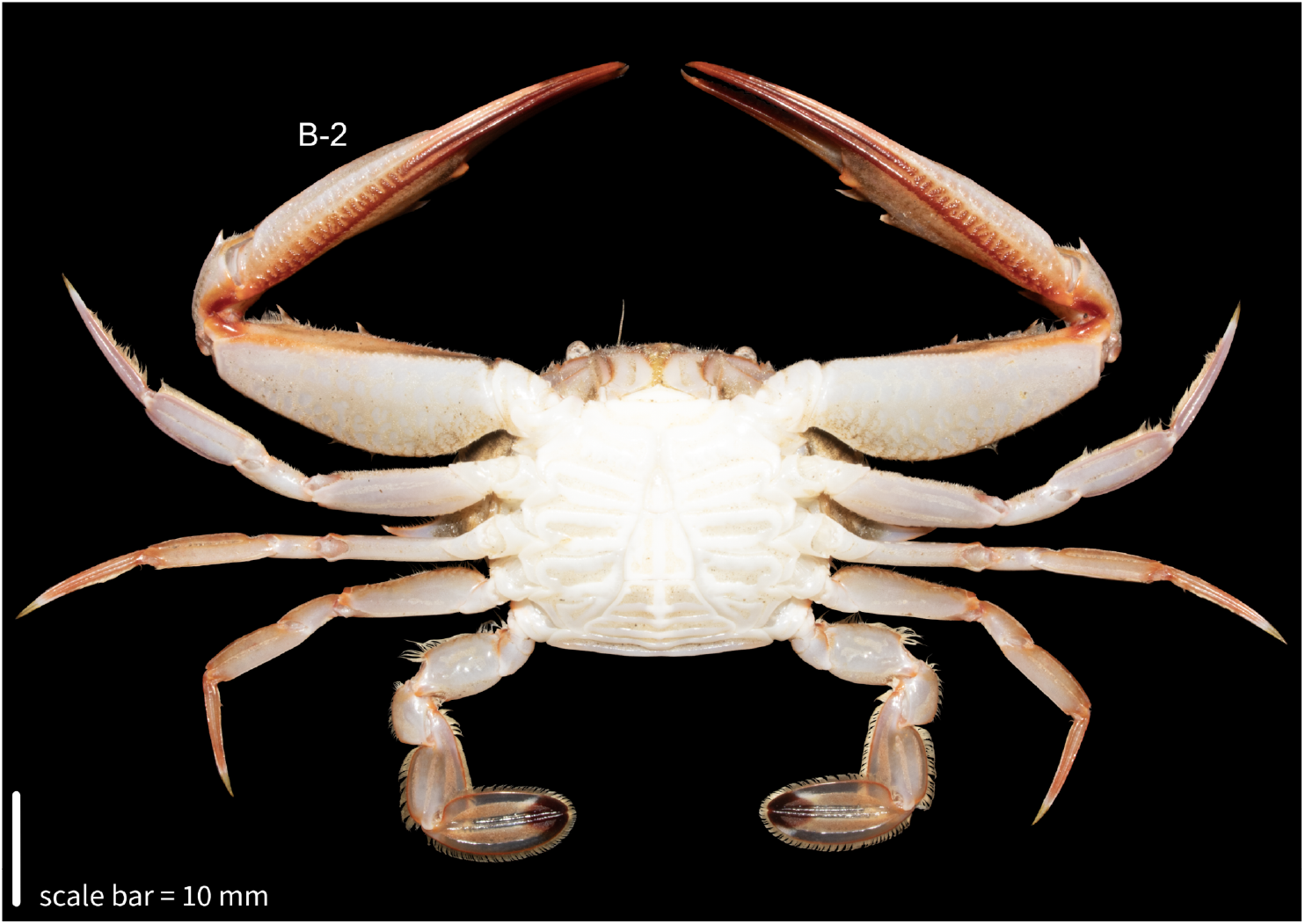
Ventral view of *Achelous hastatus* (Linnaeus, 1767) (*Decapoda*: *Brachyura*: *Portunidae*: *Achelouinae*), adult male voucher specimen (Specimen ID: IMS-AHA-001) collected from Granadilla, Tenerife, Canary Islands (28°05′06″ N, 16°29′23″ W). The specimen was encountered at a depth of 6 m in a sandy patch adjacent to mixed rocky substrate and marina walls during a night dive on 6 August 2025. Photographed under standardized laboratory conditions. Morphometric measurements: carapace width (CW) = 48.2 mm; carapace length (CL) = 23.1 mm; cheliped total length = 74.7 mm; merus length (cheliped) = 28.2 mm; carpus length (cheliped) = 12.2 mm; propodus length (cheliped) = 39.5 mm; fixed finger length = 39.5 mm; dactylus length (cheliped) = 16.8 mm; pleopod length (male) = 17.5 mm. Scale bar = 10 mm. Associated iNaturalist record: https://www.inaturalist.org/observations/304259129

### 2.3 Morphometric Measurements

Morphometric data were obtained using digital calipers with a precision of 0.1 mm. Measurements follow standardized portunid protocols as applied in comparative taxonomic studies, with terminology based on Manning & Holthuis (1981) and Spiridonov et al. (2014).

Carapace measurements:

– Carapace width (CW): maximum transverse distance across the carapace at the level of the ninth anterolateral tooth, measured perpendicular to the median longitudinal axis, including the apical tips of the ninth tooth.
– Carapace length (CL): straight-line distance from the tip of the median rostral tooth to the posterior margin of the carapace along the dorsal midline.
– Frontal width (FW): transverse distance between the inner orbital margins (postorbital lobes) at the base of the frontal region.
– Frontal spine width: transverse distance between the distal apices of the paired rostral teeth.
– Posterior width: width across the posterior carapace margin, measured between the posterolateral angles.
– Carapace height: maximum dorsal–ventral thickness of the carapace, measured perpendicular to the dorsal midline.

Cheliped measurements (left cheliped for this specimen):

– Cheliped total length: straight-line distance from the proximal margin of the coxa–basis articulation with the carapace to the tip of the fixed finger (pollex) of the propodus.
– Merus length: maximum linear distance from the proximal margin of the merus to its distal articulation with the carpus.
– Merus width: maximum transverse width of the merus, measured at its broadest point.
– Carpus length: maximum linear distance from the proximal margin to the distal articulation with the propodus.
– Carpus width: maximum transverse width of the carpus.
– Propodus length: distance from the proximal articulation with the carpus to the base of the dactylus, including the manus and fixed finger.
– Propodus width (height): maximum dorsoventral depth of the manus.
– Fixed finger length: straight-line distance from the articulation with the manus to the distal tip of the pollex.
– Dactylus length: straight-line distance from the articulation with the manus to the distal tip of the movable finger.
– Gap length: linear distance between the distal tips of the fixed finger and dactylus when fully extended

Walking leg measurements (pereiopods P2–P4, left side):

For each pereiopod, the following were measured along the maximum chord of each segment

– Merus length
– Carpus length
– Propodus length
– Dactylus length

Swimming leg measurements (pereiopod P5, left side):

– Merus length
– Carpus length
– Propodus length: chord length of the flattened paddle segment.
– Propodus width: maximum breadth of the paddle segment.
– Dactylus length: length of the terminal paddle-like segment from articulation to distal tip.

Abdominal measurements:

– Abdominal segment width (6th somite): maximum transverse width of the sixth abdominal somite, measured at its widest point.
– Telson length: distance from the anterior margin at the articulation with the sixth abdominal somite to the posterior margin of the telson.
– Pleopod length (male): maximum chord length of the first gonopod (pleopod 1) from basal articulation to distal tip.

Orbital measurements:

– Orbital width: maximum transverse width of the orbit, including the orbital margins.
– Eye diameter: horizontal diameter of the cornea at its widest point.

All measurements were recorded on the left side of the body when bilateral structures were present and are expressed in millimetres. Methodology conforms to recognized portunid morphometric standards as cited in the taxonomic literature.

### 2.4 Field Documentation and Metadata Archiving

A single adult male of *Achelous hastatus* was observed, photographed, and collected during a night dive off Tenerife. The dorsal and ventral views of the preserved voucher specimen were documented under controlled laboratory conditions (Figures B-1 and B-2). The corresponding observation was submitted to the iNaturalist platform, and associated metadata—including locality, date, depth, and photographic documentation—were archived for reference. Observation details are summarized in Table 2.

**Table 1.**

| Specimen ID | Species | Collection Locality | Coordinates | Depth (m) | Collection Date | Time (local) | Carapace Width (mm) | Carapace Length (mm) |
| --- | --- | --- | --- | --- | --- | --- | --- | --- |
| IMS-AHA-001 | *Achelous hastatus* | Granadilla, Tenerife, Canary Islands | 28°05'07"N, 16°29'23"W | 6 | 2025-08-06 | ~01:00 | 48.2 | 23.1 |

**Table 2.**
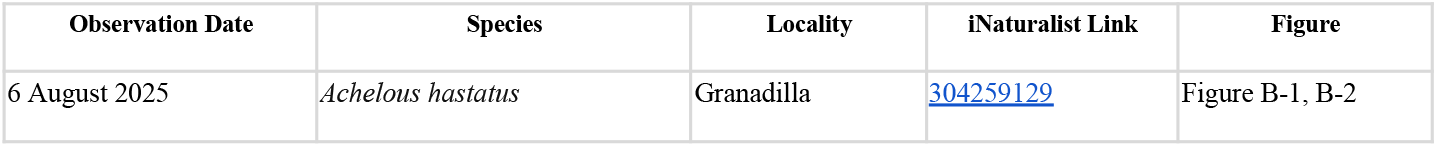

| Observation Date | Species | Locality | iNaturalist Link | Figure |
| --- | --- | --- | --- | --- |
| 6 August 2025 | *Achelous hastatus* | Granadilla | 304259129 | Figure B-1, B-2 |

### 2.5 Aggregated Occurrence Data (OBIS and GBIF)

Occurrence records for *Achelous hastatus* were retrieved from the Ocean Biodiversity Information System (OBIS) and the Global Biodiversity Information Facility (GBIF) to provide a broad-scale context for the species’ distribution.

The OBIS dataset comprised 52 georeferenced records compiled from nine contributing datasets, integrating citizen science observations, institutional collections, and historical expeditions. These included the Seawatchers Marine Citizen Science Platform (Chic & Garrabou, 2020), the Biological Reference Collections ICM CSIC (Olivas González, 2016), the Zariquiey Collection (Santos, 2016), historical decapod surveys near Alexandria, Egypt (Balss, 1936), the Mediterranean Ocean Biodiversity Information System (MedOBIS), Programa Poseidon (ECOAQUA Institute, 2015), the Aegean macrobenthic fauna dataset, the National Museum of Natural History Invertebrate Zoology Collections, and Senckenberg’s collection management system. Data were accessed via the OBIS portal (OBIS, 2025) on 11 August 2025.

The GBIF dataset comprised 958 georeferenced records worldwide, including 24 from the Canary Islands. GBIF data were accessed via occurrence download DOI https://doi.org/10.15468/dl.x5qank (GBIF.org, 2025) on 11 August 2025.

Country-level occurrence frequencies from both sources were calculated using the reported country field or deduced from coordinate-based geolocation where country attribution was missing.

## 3. Taxonomy

The examined specimen is identified as:

– ***Achelous hastatus*** (Linnaeus, 1767)
– Family: *Portunidae* Rafinesque, 1815
– Order: *Decapoda* Latreille, 1802
– Class: *Malacostraca* Latreille, 1802

The full classification is provided below, based on DecaNet and the World Register of Marine Species (WoRMS):

Kingdom ***Animalia*** Linnaeus, 1758

Phylum ***Arthropoda*** Latreille, 1829

Subphylum ***Crustacea*** Brünnich, 1772

Superclass ***Multicrustacea*** Regier et al., 2010

Class ***Malacostraca*** Latreille, 1802

Subclass ***Eumalacostraca*** Grobben, 1892

Superorder ***Eucarida*** Calman, 1904

Order ***Decapoda*** Latreille, 1802

Suborder ***Pleocyemata*** Burkenroad, 1963

Infraorder ***Brachyura*** Linnaeus, 1758

Section ***Eubrachyura*** De Saint Laurent, 1980

Subsection ***Heterotremata*** Guinot, 1977

Superfamily ***Portunoidea*** Rafinesque, 1815

Family ***Portunidae*** Rafinesque, 1815

Subfamily ***Achelouinae*** Spiridonov, 2020

Genus ***Achelous*** De Haan, 1833

Species ***Achelous hastatus*** (Linnaeus, 1767)

*Achelous hastatus* (Linnaeus, 1767) is currently regarded as an accepted taxon in WoRMS (AphiaID: 1474844). The species was originally described as *Cancer hastatus* Linnaeus, 1767, and has subsequently appeared in the literature under several superseded combinations and junior subjective synonyms, including *Portunus hastatus* (Linnaeus, 1767), *Neptunus hastatus* (Linnaeus, 1767), and *Lupa Dufourii* Desmarest, 1825. It belongs to the subfamily *Achelouinae* within *Portunidae*, a group morphologically characterized by a generally hexagonal carapace with nine anterolateral teeth, the ninth tooth being markedly elongate, and by the presence of natatory fifth pereiopods adapted for swimming. *Achelous hastatus* is a widely distributed portunid crab occurring in marine, brackish, and occasionally freshwater environments, and is represented in both recent and fossil records (DecaNet eds., 2025).

## 4. Results

A single adult male specimen of Achelous hastatus (Linnaeus, 1767) was collected on 6 August 2025 from a sandy patch adjacent to mixed rocky substrate and marina walls at 6 m depth off Granadilla, Tenerife, Canary Islands. The specimen was photographed in dorsal and ventral views under controlled lighting conditions in a laboratory setting. Two high-resolution images (Figures B–1 and B–2) were produced to illustrate the general morphology.

Morphometric measurements of the examined specimen are summarised in Table 3. Measurement abbreviations correspond to the definitions provided in the Material and Methods section.

**Table 3.**
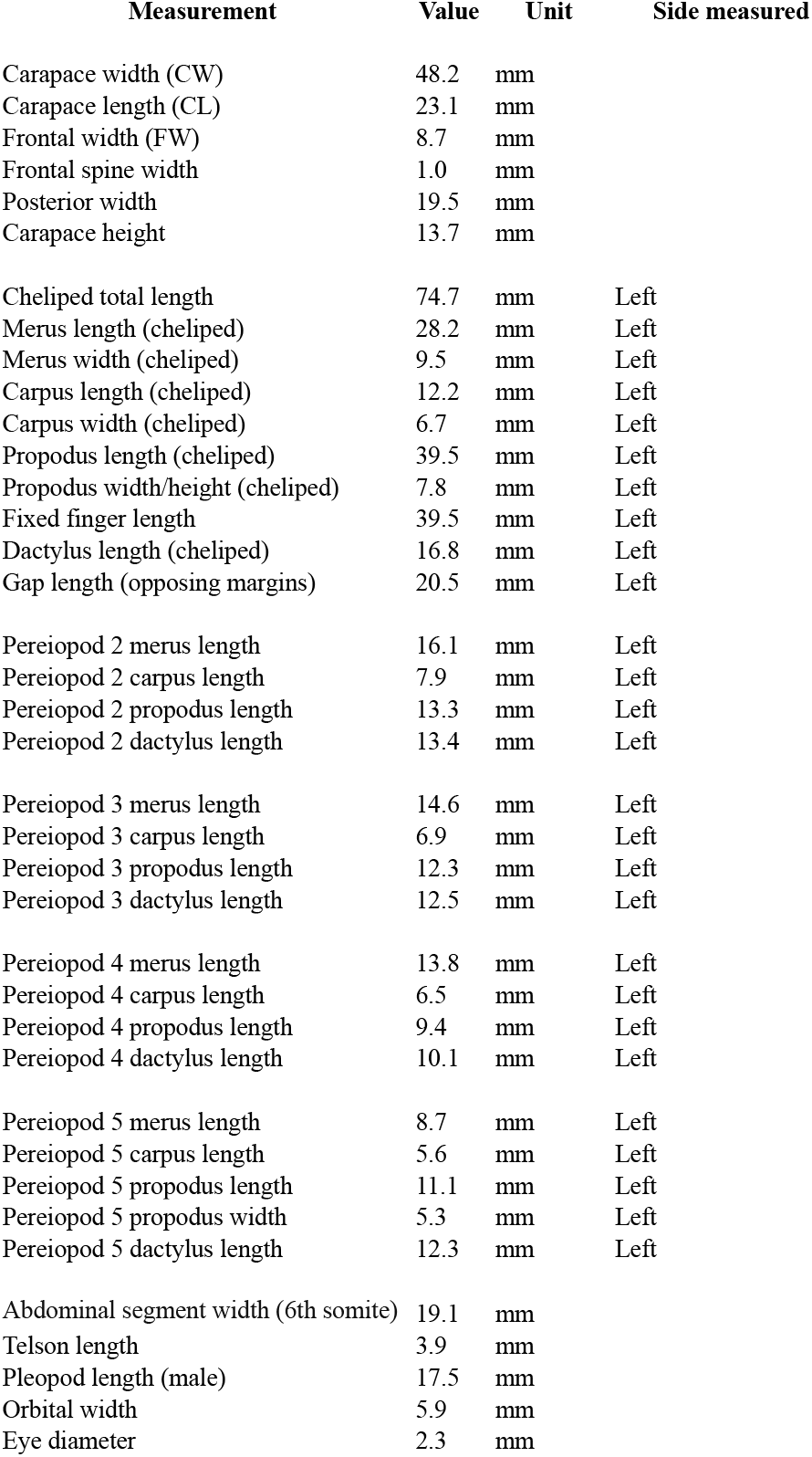

| Measurement | Value | Unit | Side measured |
| --- | --- | --- | --- |
| Carapace width (CW) | 48.2 | mm |  |
| Carapace length (CL) | 23.1 | mm |  |
| Frontal width (FW) | 8.7 | mm |  |
| Frontal spine width | 1.0 | mm |  |
| Posterior width | 19.5 | mm |  |
| Carapace height | 13.7 | mm |  |
| Cheliped total length | 74.7 | mm | Left |
| Merus length (cheliped) | 28.2 | mm | Left |
| Merus width (cheliped) | 9.5 | mm | Left |
| Carpus length (cheliped) | 12.2 | mm | Left |
| Carpus width (cheliped) | 6.7 | mm | Left |
| Propodus length (cheliped) | 39.5 | mm | Left |
| Propodus width/height (cheliped) | 7.8 | mm | Left |
| Fixed finger length | 39.5 | mm | Left |
| Dactylus length (cheliped) | 16.8 | mm | Left |
| Gap length (opposing margins) | 20.5 | mm | Left |
| Pereiopod 2 merus length | 16.1 | mm | Left |
| Pereiopod 2 carpus length | 7.9 | mm | Left |
| Pereiopod 2 propodus length | 13.3 | mm | Left |
| Pereiopod 2 dactylus length | 13.4 | mm | Left |
| Pereiopod 3 merus length | 14.6 | mm | Left |
| Pereiopod 3 carpus length | 6.9 | mm | Left |
| Pereiopod 3 propodus length | 12.3 | mm | Left |
| Pereiopod 3 dactylus length | 12.5 | mm | Left |
| Pereiopod 4 merus length | 13.8 | mm | Left |
| Pereiopod 4 carpus length | 6.5 | mm | Left |
| Pereiopod 4 propodus length | 9.4 | mm | Left |
| Pereiopod 4 dactylus length | 10.1 | mm | Left |
| Pereiopod 5 merus length | 8.7 | mm | Left |
| Pereiopod 5 carpus length | 5.6 | mm | Left |
| Pereiopod 5 propodus length | 11.1 | mm | Left |
| Pereiopod 5 propodus width | 5.3 | mm | Left |
| Pereiopod 5 dactylus length | 12.3 | mm | Left |
| Abdominal segment width (6th somite) | 19.1 | mm |  |
| Telson length | 3.9 | mm |  |
| Pleopod length (male) | 17.5 | mm |  |
| Orbital width | 5.9 | mm |  |
| Eye diameter | 2.3 | mm |  |

The observed morphological characters correspond closely with the diagnostic description of Achelous hastatus provided by Rodrigues et al. (2017). These include a broadly hexagonal carapace with nine distinct anterolateral teeth, the ninth markedly elongate, acute, and projecting laterally beyond the orbital margin; a frontal margin divided into four distinct lobes; robust chelipeds with a proportionally narrow manus and comparatively slender fixed and movable fingers; and a fifth pereiopod bearing an ovate, setose natatory dactylus. The male abdomen is narrow, with somites 3–5 fused and the telson triangular, consistent with the species-level diagnosis for A. hastatus.

## 5. Discussion

The present record of Achelous hastatus from Granadilla, Tenerife, adds voucher-backed, morphometrically documented evidence of the species in the Canary Islands. While the species is well represented in occurrence databases such as GBIF and OBIS (GBIF.org, 2025; OBIS, 2025), regional records often lack detailed morphometric datasets or high-resolution imaging. Our specimen, preserved with associated dorsal and ventral photographs, provides a verifiable reference point for future taxonomic and ecological studies in the Macaronesian region.

The observed morphological characters, including the nine anterolateral teeth with an elongate ninth, four-lobed frontal margin, and ovate setose natatory dactylus, are fully consistent with the diagnostic criteria outlined by Rodrigues et al. (2017) for A. hastatus. This agreement underscores the stability of these traits across the species’ range and supports the accuracy of regional identifications when based on high-quality imagery and morphometric confirmation.

Ecologically, the specimen was observed at night within a sandy patch adjacent to mixed rocky substrate, in association with a feeding aggregation of at least six conspecifics actively scavenging on organic remains. This aligns with prior observations of opportunistic feeding and nocturnal foraging behaviour in portunid crabs (Spiridonov et al., 2014), but direct documentation of group scavenging on vertebrate remains in A. hastatus appears rare in the literature. Such behaviour may reflect opportunistic exploitation of anthropogenic food inputs in nearshore environments.

The Canary Islands lie near the north-eastern limit of the species’ confirmed Atlantic–Mediterranean distribution. The occurrence data synthesised from OBIS (2025) show 52 georeferenced records compiled from nine datasets, with Spain (including the Canary Islands) accounting for the majority. GBIF data further emphasise the species’ dominance in Spanish waters, with 657 of 958 global records originating from Spain (GBIF.org, 2025). This congruence between datasets supports the species’ established status in the region.

By contributing a well-documented, geo-referenced voucher specimen with precise morphometrics, this study strengthens the biogeographic dataset for A. hastatus and highlights the value of combining field observation, specimen preservation, and open-access biodiversity databases in marine faunal research.

## Notes

### Competing Interest Statement

The authors have declared no competing interest.

